# Spatial chromosome organization and adaptation of *Escherichia coli* under heat stress

**DOI:** 10.1101/2024.05.31.596820

**Authors:** Xu-Ting Wang, Bin-Guang Ma

## Abstract

The spatial organization of bacterial chromosomes is crucial for cellular functions. It remains unclear how bacterial chromosomes adapt to high temperature stress. This study delves into the 3D genome architecture and transcriptomic responses of *Escherichia coli* under heat stress condition to unravel the intricate interplay between chromosome structure and environmental cues. By examining the role of macrodomains, chromosome interaction domains (CIDs), and nucleoid-associated proteins (NAPs), this work unveils the dynamic changes in chromosome conformation and gene expression patterns induced by high temperature stress. It was observed that under heat stress, short-range interaction frequency of chromosome decreased, while the long-range interaction frequency of the Ter macrodomain increased. Furthermore, two metrics, namely, Global Compactness (GC) and Local Compactness (LC), were devised to measure and compare the compactness of chromosomes based on their 3D structure models. The findings in this work shed light on the molecular mechanisms underlying thermal adaptation and chromosomal organization in bacterial cells, offering valuable insights into the complex interrelationships between environmental stimuli and genomic responses.

## 1. Introduction

In cellular organisms, chromosomes undergo significant compression to fit within the confines of the cell. The mechanisms by which organisms achieve this compression while still enabling vital processes like replication and transcription remain elusive. The development of chromosome conformation capture (3C) technologies has provided crucial support for analyzing chromosome organization *in vivo* [1]. Unlike eukaryotes, bacteria lack a nucleus encased in a nuclear membrane, resulting in DNA organization into nucleoids. Recently, there has been a growing application of 3C technologies to investigate the structural organization of bacterial nucleoids [2–4]. Macrodomain organization is a prevalent feature across bacterial nucleoids [5–7]. The Ori and Ter macrodomains in *E. coli* were initially identified using fluorescence in situ hybridization (FISH) [5], and their uneven distribution within the circular chromosome was directly visualized through fluorescence microscopy [8]. Global 3C data have validated the presence of four macrodomains and two non-structural regions in logarithmically growing cells [9, 10]. Macrodomain formation is believed to be closely linked to Structural Maintenance of Chromosomes (SMC) proteins. Homologs of SMC proteins in *E. coli* are MukBEF proteins, which are implicated in chromosome condensation over long distance extending beyond the Ter macrodomain [11].

3C coupled with sequencing (3C-seq) data analysis has revealed the presence of 31 chromosome interaction domains (CID) in *E. coli* [9]. It is hypothesized that the macrodomains of *E. coli* may be comprised of nested CIDs, with a significantly higher frequency of interactions occurring within CIDs compared to interactions between them [12]. Some studies suggest that CID formation in *E. coli* may be stochastic and subject to variation under different growth conditions [13], with the underlying mechanism remaining elusive. It is speculated that CID formation may be influenced by factors such as DNA supercoiling, macromolecular crowding, and transcription processes. Nucleoid-associated proteins (NAPs) play a pivotal role in organizing the nucleoid structure of *E. coli*, with 12 common NAPs identified, including Fis, HU, H-NS, INF, and StpA [14]. Super-resolution fluorescence microscopy reveals their distinct localization patterns: H-NS exhibits a clustered distribution, while HU, Fis, IHF and StpA are dispersed throughout the nucleoid [15]. 3C data indicate that H-NS predominantly regulates short-distance interactions, whereas HU and Fis modulate long-distance interactions through distinct mechanisms [9]. Additionally, the interactions between NAPs and DNA, alongside the concentration of metabolites, proteins, water, and salt in the cytoplasm, influence the solvent property of the cytoplasm, thereby impacting nucleoid spatial organization [16–21].

The heat stress response in *E. coli* has garnered significant attention since its inception [22–26]. Traditional research methodologies have elucidated numerous molecular mechanisms underlying heat response regulation [26–29], and these findings have found broad application, particularly in synthetic biology [25, 30]. A previous investigation has identified a correlation between the expression levels of heat stress-responsive genes and their sequence properties [28]. The supercoil structure of bacterial DNA is influenced by transcription levels [31], and changes in temperature affect transcription levels and temperature-sensitive mRNA conformation [26]. However, it remains unclear how temperature alterations induce changes in the chromosome structure, potentially affecting the expression levels of NAPs, and how the variations of NAPs expression levels reciprocally influence chromosome conformation in *E coli*. To address these questions, we need to study the 3D genome under heat stress condition.

In this study, we employed 3C-seq technology to elucidate the chromosome conformations of *E. coli* under normal temperature (37°C) and high temperature (45°C) conditions. Additionally, RNA-seq technology was adopted to assess the transcription levels of *E. coli* genes across varying temperatures. Through integration of 3D genomic and transcriptomic analyses, we observed significant alterations in both transcription levels and the 3D organization of the *E. coli* nucleoid in response to high temperature stress. Specifically, we observed a decrease in the interaction frequency between DNA segments within a range of approximately 200 kb, accompanied by a relative increase in the interaction frequency among DNA segments spanning over 1000 kb in the high temperature environment. Furthermore, based on 3D structural models, analysis of Global Compactness (GC) reveals a trend toward increased chromosome compaction under high temperature stress, while Local Compactness (LC) exhibits distinct changes across different chromosome regions. Through comprehensive analysis of gene transcription levels and chromosome structural features, we uncovered a close association between alterations in chromosome organization and changes in transcription level induced by high temperature. These findings underscore the intricate interplay between chromosomal architecture and gene expression in response to environmental stress.

## 2. Materials and methods

### 2.1. Bacterial culture and heat treatment

The bacterial strain used in this study was *Escherichia coli* MG1655 preserved in our laboratory. LB medium was used for cultivating *E. coli* in the experiment. *E. coli* cells stored at -80 ℃ were inoculated in fresh LB medium at 1:50 ratio before culturing, then cultured at 37 ℃ overnight, and then inoculated in LB medium at 1:100 ratio for culturing at 37 ℃. When the OD_600_ of culture medium reached 2 (about 1.5 h), the culture medium was divided into two groups: normal temperature group and high temperature group. The normal temperature group was continued at 37 ℃, and the culture temperature in the high temperature group was raised to 45 ℃ (by placing the culture medium in a water bath of 50 ℃ for 1 min and 20 s) to continue. At two time points (namely, 10 min and 2.5 h after heat treatment), 1 mL of the culture was sampled in the normal temperature group and the high temperature group, respectively, to prepare 3C library, RNA-seq library and nucleoid fluorescence imaging experiments. The regents used in the experiments are listed in **Table S1**.

### 2.2. 3C and RNA sequencing

The method described in Lioy and co-authors’ work [9] was modified to generate the 3C library of *E. coli*. The *E. coli* cells were cultured at 45℃ for 10 minutes (thermal logarithmic growth phase, simplified as: Therm_Log) and 2.5 hours (thermal stationary growth phase, simplified as: Therm_Sta) and at 37℃ for the same time points (Norm_Log and Norm_Sta) in four groups of culture media, and 1mL of culture media in each group was sampled for the construction of 3C and RNA sequencing library, respectively. The detailed process for 3C and RNA library construction is described in **Supplementary Methods section 2.1**. The resulting 3C and RNA libraries were sequenced with Illumina HiSeq 2500 equipment (150 bp pair-end sequencing).

### 2.3. Fluorescence imaging of bacterial nucleoid

The *E. coli* culture medium was centrifuged at 3500 × g for 10 minutes to remove the supernatant, and then 1 × DAPI staining solution was added to re-suspend the bacteria. After incubation at room temperature for 30 minutes, the supernatant was centrifuged again, washed twice with 1 × PBS, and re-suspended with 1 × PBS. Using Nikon Ti2-E microscope for multi-channel (bright field DIA and dark field DAPI) imaging, the images of the two channels could be superimposed to obtain the contours of *E. coli* cells and their nucleoids. OpenCV was used for image recognition, and the length and width of the outer contour of the bacteria nucleoid in each fluorescence image were recorded.

### 2.4. Interaction matrix and CID analysis

The procedure in Lioy and co-authors’ work [9] was used to generate chromosome interaction matrices at 5 kb resolution (bin size) based on the data obtained from sequencing of the 3C library. The generated interaction matrix was normalized by SCN method [32] to facilitate subsequent comparative analysis. The short-range interaction frequency and proportion were calculated to reflect how a target bin interacts with its neighboring bins on the genome (see **Supplementary Methods section 2.2 for details**). According to the definition in Dixon and co-authors’ work [33], the directionality index (DI) was calculated for each interaction matrix by using the script of Lioy et al. [9]. According to definition, DI reflects the tendency of each bin to interact with the upstream and downstream bins along the chromosome direction. Therefore, the alternating position of positive and negative DI value is considered as the location of CID boundary.

### 2.5. Transcription level and correlation analysis

After removing the adapter from the Illumina sequencing data, the software Cutadapt was used to remove the bases of reads with low quality at the head and tail. Sequencing data were aligned to the reference genome (U000913.3) using the software TopHat calling Bowtie2. The FPKM values for each gene were calculated by combining the results of the comparison with the reference annotation information using the software Cufflinks. Since the interaction frequency data are in matrix form, with segment of 5 kb size as units, and the transcription levels are in gene units with large fluctuations, it is impossible to directly analyze the correlation. Firstly, we divided the transcription level data into 5 kb segment (bin) according to the location of gene, counted the transcription level of each bin, and then calculated the log2 value of this level to make it relatively centralized and convenient for comparison. Then, the values of the diagonal in the interaction matrix were used to calculate the Z-score of interaction frequency, and this Z-score was used for Pearson correlation analysis with the Z-score of transcription level. The software tools used in this study are summarized in **Table S2**.

### 2.6. Chromosome 3D model construction and compactness calculation

Using the EVR program [34], the normalized interaction matrices were converted to 3D structure models. The generated models could be directly visualized using PyMol. To facilitate comparison between 3D models, we devised two indicators for measuring the compactness of 3D chromosome structure: Global Compactness (GC) and Local Compactness (LC).

GC is defined as **Eq. 1**:

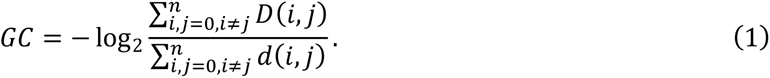

In this formula, GC is the Global Compactness, *i* and *j* are indexes for bins, *D* (*i*, *j*) is the Euclidean distance between bin *i* and bin *j* in the chromosome 3D structure model, and *d* (*i*, *j*) is the distance between bin *i* and bin *j* on a reconstructed circle. The details for the calculation of GC are described in **Supplementary Methods section 2.3**.

LC is defined as **Eq. 2**:

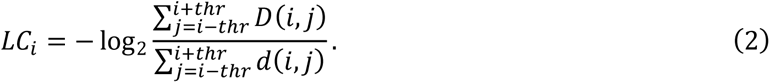

In this formula, *LC_i_* is the local compactness of bin *i*, *D* (*i*, *j*) is the Euclidean distance between bin *i* and bin *j* in the chromosome 3D structure model, *d* (*i*, *j*) is the distance between bin *i* and bin *j* on a reconstructed straight line within the local range of bin *i*, and *thr* is the threshold for local range definition. The larger the *LC_i_*, the higher the chromosome compactness at the location of bin *i*. The details for the calculation of LC are described in **Supplementary Methods section 2.4**.

## 3. Results

### 3.1. Chromosome interaction decreases with linear genomic distance in all conditions

For the four groups (Norm_Log, Norm_Sta, Therm_Log, Therm_Sta) of *E. coli* culture samples, 3C-seq experiments were conducted, yielding interaction frequencies between chromosomal DNA fragments, which reflect the spatial organization of *E. coli* chromosome under these conditions. Analysis involving the linear genomic distance and spatial interaction frequency of DNA segments revealed notable trends. **Figure1A** illustrates that the spatial interaction frequency between DNA segments inversely correlates with linear genomic distance under all growth conditions. Most chromosome interactions occur within a linear distance of 500 kb, and beyond this range, the interaction frequencies decline to very low levels. Upon closer examination of the interaction frequency patterns within 500 kb, we observed that samples from Norm_Log group exhibit the highest interaction frequencies at equivalent linear distances, and samples from the Norm_Sta group exhibit the lowest interaction frequencies, whereas samples from the Therm_Log and Therm_Sta groups display interaction frequencies of intermediate levels.

### 3.2. The long-range (>200 kb) interaction of Ter macrodomain increases under the heat stress

We calculated the ratio of the interaction matrix under heat stress and the interaction matrix under normal temperature, and drew the heat maps (**Figure 1B**), from which the change of *E. coli* chromosome interaction under heat stress can be obviously observed. As shown, compared with the normal temperature condition, the interaction frequencies of logarithmic and stationary phases under heat stress condition increased relatively in the range above 1000kb, while the reduction of interaction frequency was mainly concentrated in the range below 1000kb, as shown by the black dashed lines in **Figure 1B**. The lower two subfigures of **Figure 1B** show the ratio of the interaction frequency over a range of 1000 kb, where the green dashed lines represent 200 kb. As shown in the left subfigure (corresponding to Therm_Log condition), the DNA interaction frequency within 200 kb is apparently reduced. In the stationary phase under heat stress (right subfigure), the interaction frequency of Ter macrodomain is slightly reduced in the range of 200 kb but apparently increased beyond, meaning that more extensive interactions occur between Ter macrodomain and the DNA segments beyond a linear genomic distance of 200kb.

**Figure 1.**
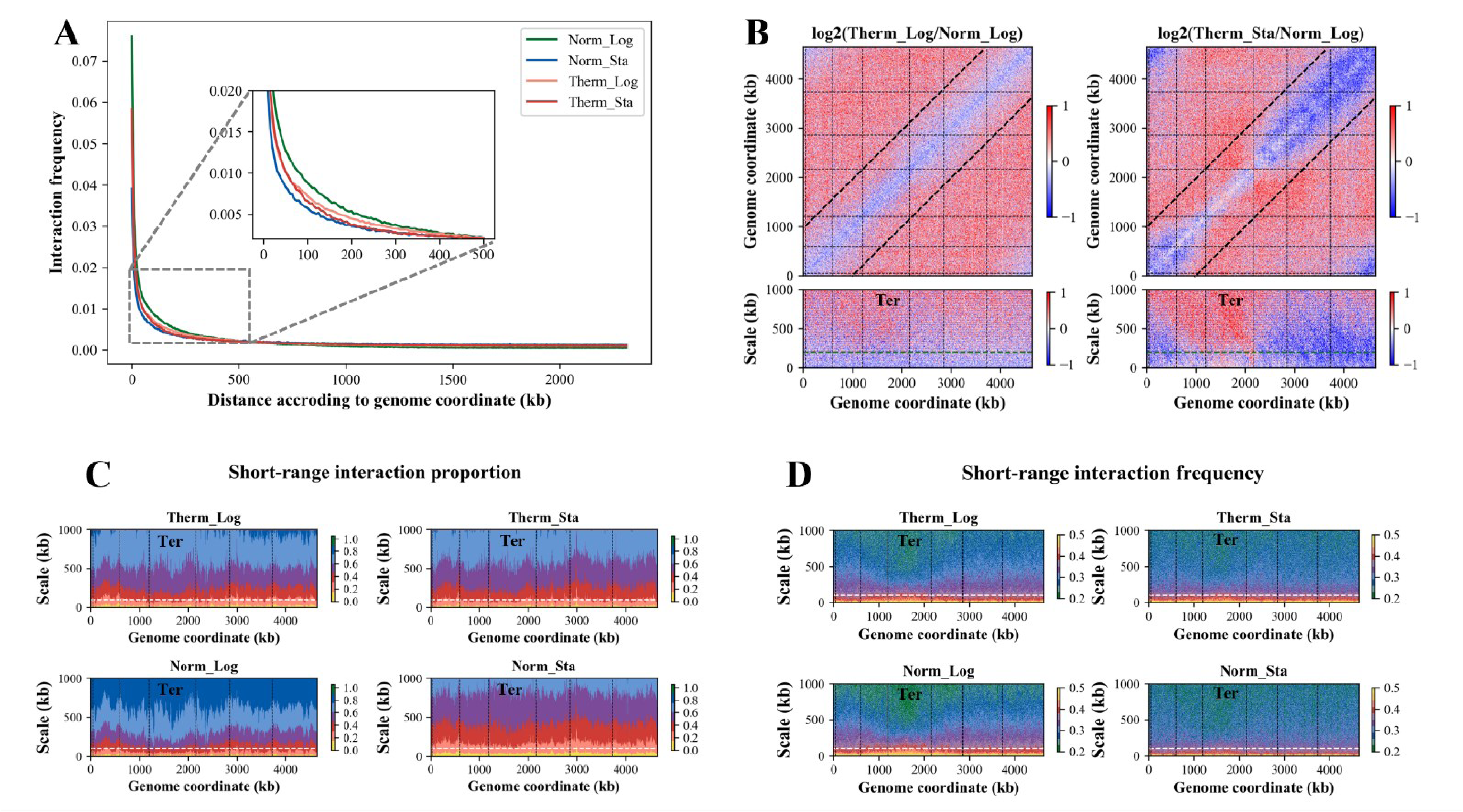
*E. coli* chromosome interactions under different growth conditions. (A) Interaction frequency of DNA segments varies with the linear genomic distance. (B) Heat maps for the ratio of interaction frequency between high and normal temperature growth conditions. Blue indicates a decrease in interaction frequency under high temperature condition, and red indicates an increase. The green dashed lines on the figure indicate 200 kb. (C) Short-range (<100 kb) interaction proportions under different growth conditions. (D) Short-range (<100 kb) interaction frequencies under different growth conditions. The white dashed lines in the subfigures C and D indicate 100 kb. The black dotted vertical lines in the subfigures B, C, D indicate the boundaries of macrodomains.

### 3.3. Short-range (<100 kb) interactions are significantly reduced under the heat stress

We investigated the effect of heat stress on short-range (<100 kb) interactions within the *E. coli* genome. The short-range chromosome interactions are characterized by analyzing the frequency and proportion of interactions between DNA segments in proximity. Interaction frequency describes the average number of interactions between a target bin and its flanking bins (upstream and downstream) at a specific distance. The interaction proportion, on the other hand, quantifies the cumulative interaction frequency within a defined window around the target bin. These two parameters capture distinct aspects of short-range interactions.

Our analysis revealed that a significant proportion (>20%) of all interactions occur within a 100 kb window, which represents approximately 2.5% of the genome length (**Figure 1C**). The interaction ratio further highlights that over 60% of the target bin’s interactions originate from bins within a 500 kb window (**Figure 1C**). These findings suggest that chromosome interactions are primarily confined to a range of <500 kb linear genomic distance. To further explore these short-range interactions, we calculated the interaction frequency within a 1 Mb window surrounding each bin. The resulting maps (**Figure 1D**) clearly demonstrate a reduced interaction frequency within a range of 100 kb. Interestingly, the Ter macrodomain exhibits a distinct interaction profile. Within 100 kb, the Ter macrodomain displays a higher interaction frequency compared to other regions. However, beyond 100 kb, the interaction frequency of the Ter macrodomain drops significantly, suggesting a potential role for MatP in establishing domain-specific interaction insulation, in line with previous reports [35, 36].

### 3.4. Transcription level positively correlates with interaction frequency

We explored the relationship between chromosome interaction and gene expression in *E. coli* under different growth conditions. To analyze the correlation between DNA interaction frequency and transcription level, we assigned the transcription levels of genes into 5 kb bins (see **Methods** for details). Then, the average transcription level of each bin and the Z-score of the interaction frequency in the upstream and downstream 10 kb range of the bin were calculated. The Z-score curves of interaction frequency and the corresponding transcription levels are shown in **Figure 2A** for the four growth conditions, respectively. As can be seen, there are clear positive correlations between the trends of interaction frequency and transcription level for the samples from these conditions. The correlation coefficients for the two log-phase samples are greater than 0.5, and the correlation coefficients for the two stationary-phase samples are around 0.4, and all of them are highly significant (*p*-value near 0 for each sample). Changes in growth conditions affect the interaction frequency and transcription level of DNA, while the correlation between them remains positive under all conditions.

**Figure 2.**
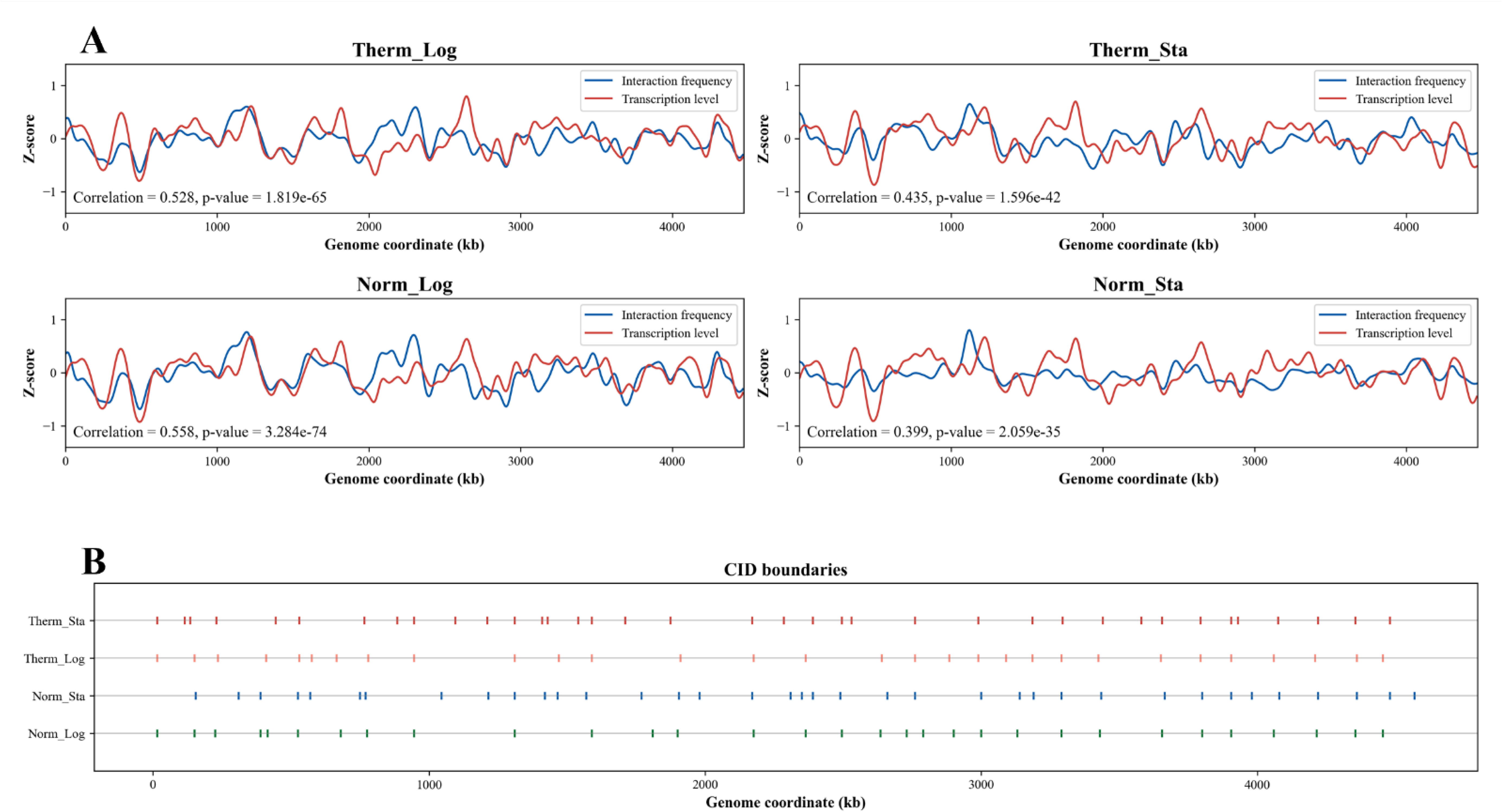
The relationship between DNA interaction frequency and transcription level and the CID boundaries under different growth conditions. (**A**) Correlation between interaction frequency and transcription level of DNA segments under different growth conditions. Blue lines represent the Z-score of DNA interaction frequency, and red lines represent the Z-score of DNA transcription level. The lower left corner of each subfigure displays the correlation coefficient and corresponding significance level (*p*-value). (B) CID boundaries under different growth conditions. The horizontal lines represent the genomic coordinates, and the points on each line represent the CID boundaries.

### 3.5. Differential CID boundary genes are related with cell wall and membrane

We identified Chromosome Interaction Domains (CIDs) in the *E. coli* genome by calculating directionality index (DI) [33]. For the four growth conditions, the distribution of CID boundaries across the genome for each condition is depicted in **Figure 2B**. Interestingly, while the total number of CIDs remains relatively constant between normal and high temperature logarithmic phases (31 vs. 30), their boundary locations differ. Gene Ontology (GO) enrichment analysis of the genes located at these differential boundaries revealed a significant enrichment for terms related to lipopolysaccharide (LPS) and oligosaccharides (**Figure S3**). As LPS is a crucial component of the *E. coli* cell wall, this result may suggest that heat stress alters LPS structure, triggering a signaling cascade that activates various cellular stress responses. The analysis of the stationary phases identified 37 CIDs in both temperature conditions. Notably, the stationary phases display a higher density of CID boundaries around the Ter macrodomain compared to the logarithmic phases, accompanied by a general decrease in CID size. Furthermore, GO enrichment analysis of the genes located at the differential CID boundaries in the high temperature stationary phase identified an enrichment for terms associated with transmembrane transport (**Figure S3**). This observation is consistent with the notion that heat stress can damage the cell wall, necessitating enhanced transmembrane transport activity for intracellular repair processes. These findings collectively support a strong link between changes in growth conditions and the dynamic reorganization of the *E. coli* chromosome structure.

### 3.6. The Global Compactness of *E. coli* chromosome is higher in stationary phase and high temperature environment

To further explore chromosome changes, we employed the EVR program [34] to generate 3D structure models of the *E. coli* chromosome under various growth conditions (**Figure 3A**). The models depict four macrodomains (Ori, Ter, Left, Right) and two non-structured regions. All models share a crescent-shaped ring structure with numerous folds. Notably, Ori and Ter occupy opposite ends of the models, with Left and Right macrodomains flanking Ter but separated without significant intertwining.

**Figure 3.**
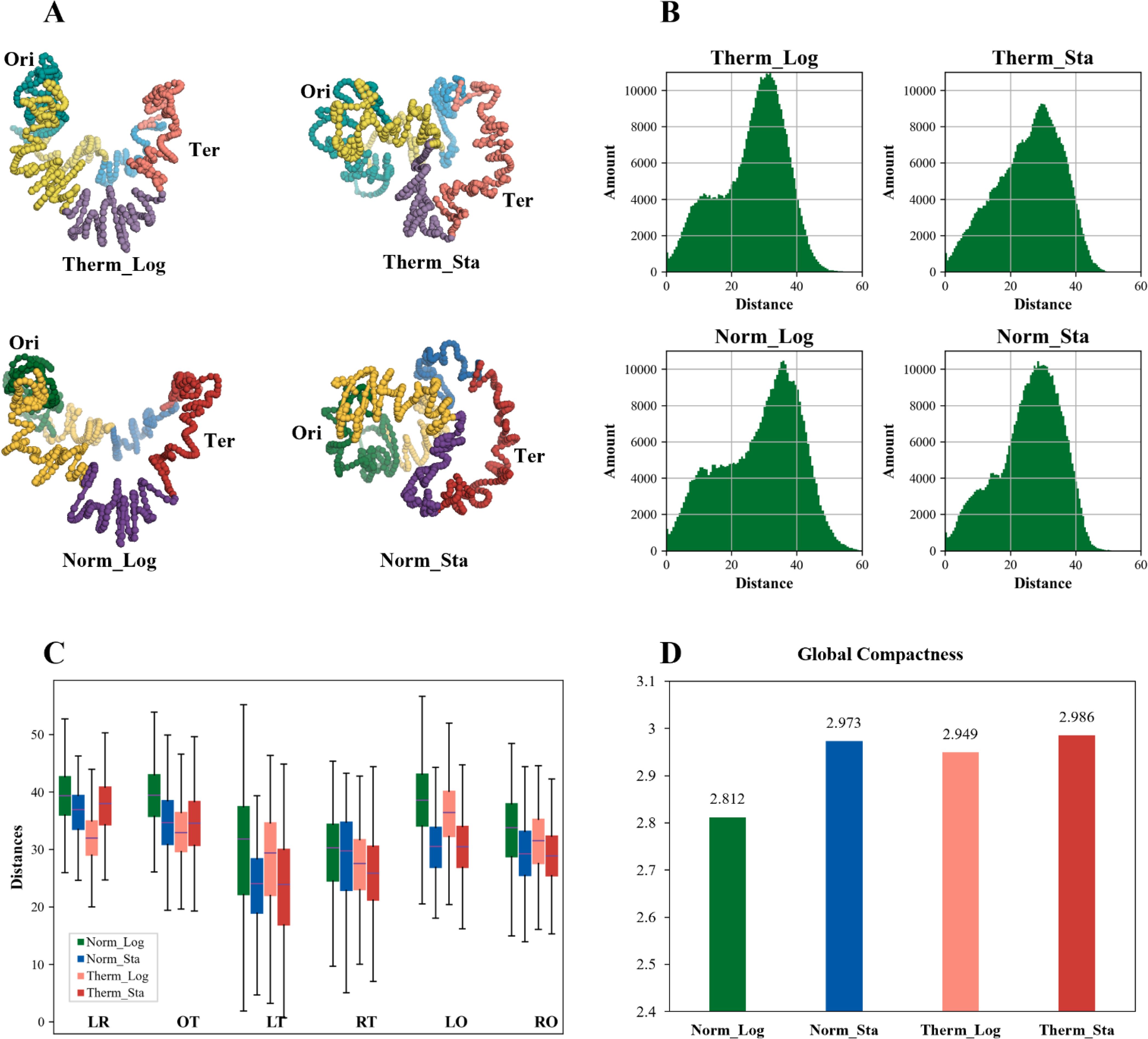
The 3D structural features of *E. coli* chromosome under different growth conditions. (**A**) The 3D structural models of *E. coli* chromosome under different growth conditions. In the Norm_Log and Norm_Sta models, green is the Ori macrodomain, red is the Ter macrodomain, purple and blue are the Left and Right macrodomains, respectively, and yellow is the non-structured regions; in the Therm_Log and Therm_Sta models, lighter colors are used correspondingly. (B) Distribution of distance between points (bins) in these 3D models. (C) Spatial distances between bins of macrodomains in the 3D models of *E. coli* chromosome under different growth conditions. O: Ori macrodomain; T: Ter macrodomain, L: Left macrodomain; R: Right macrodomain. OT represents the distances between the bins in the Ori macrodomain and the bins in the Ter macrodomain, and so on. (D) Global Compactness of the *E. coli* chromosome models under different growth conditions.

While all models exhibit a crescent-shaped ring, subtle differences exist upon closer inspection. The log-phase models appear highly similar, although the Therm_Log model displays a slightly closer proximity between Ori and Ter. The stationary-phase models appear smaller than their log-phase counterparts, with a significantly reduced distance between Ori and Ter. To quantify these observations, we analyzed the distance distribution for each point (bin) pair within the 3D models **(Figure 3B**). The log-phase models exhibit a near-bimodal distribution, with a dominant peak around 30-40 units and a less prominent peak near 10 units. In contrast, the stationary-phase models display a single peak at 30-40 units. This suggests a decrease in the frequency of short distance (around 10 units) and an increase in the frequency of intermediate distance (around 20 units) in the stationary-phase models, particularly under heat stress. To further substantiate these findings, we analyzed distance variations between macrodomain bins (namely, the bins belonging to each macrodomain, respectively) within the models (**Figure 3C**). The data reveal a consistent trend: distances between any two macrodomains tend to be smaller in the high temperature log-phase models compared to the normal temperature log-phase models. Additionally, distances between bins within each macrodomain also shows a decrease trend for these macrodomains (data not shown), suggesting a genome-wide trend of chromosome condensation under heat stress.

To quantify the overall chromosome compactness, we defined a metric termed Global Compactness (GC). GC calculation involves reconstructing the chromosome model into a circle and then calculating the negative logarithm of the ratio between the sum of distances in the original model and the sum of distances in the reconstructed circle (see **Methods** and **Supplementary Methods** for details). Higher GC values indicate greater compactness or condensation. GC values for the Norm_Log and Norm_Sta models were 2.812 and 2.973, respectively, and 2.949 and 2.986 for the Therm_Log and Therm_Sta models, respectively (**Figure 3D**). These results support the observation that chromosomes under high temperature are more compact than those under normal temperature at the same growth stage. Additionally, the degree of chromosome condensation is higher in the stationary phase compared to the logarithmic phase under the same temperature condition. This aligns with previous studies employing sucrose density gradient centrifugation to isolate *E. coli* nucleoids at different growth stages, which reported a more compact nucleoid structure in the stationary phase [37]. In conclusion, both transition to stationary phase and exposure to heat stress promote a higher degree of overall chromosome condensation in *E. coli*.

### 3.7. No simple relationship between chromosome Global Compactness and the nucleoid size of *E. coli* cell

*E. coli* typically appears as a short, straight rod-shaped bacterium. However, high temperature stress can influence cell size. Previous studies [38, 39] reported an increase in *E. coli* cell length at elevated temperatures (up to 50°C). Consistent with these reports, DAPI staining and fluorescence microscopy revealed that *E. coli* in the high temperature environment displayed a changed morphology (**Figure 4A**). In order to accurately compare the size variations of *E. coli* nucleoid under different growth conditions, we utilized microscopic image analysis to measure the width and length of each nucleoid. Approximately 2000 *E. coli* cells were analyzed for each growth condition. The statistical result in **Figure 4B** indicates that the widths of *E. coli* nucleoid do not exhibit significant change between the logarithmic and stationary phases at normal temperature. Conversely, in a high temperature environment, the widths of *E. coli* nucleoid decrease significantly for both the logarithmic and stationary phases, with significant higher values in stationary phase than in logarithmic phase. The statistical results for length are presented in **Figure 4C**, revealing that compared to the logarithmic phase, the lengths of *E. coli* nucleoid are notably shorter during the stationary phase at normal temperature. However, there is no significant difference in length for the logarithmic phase between high and normal temperatures, and meanwhile the nucleoid lengths are significantly larger in high temperature stationary phase, compared to other three phases. The 3D genomic data demonstrate that *E. coli* chromosomes exhibit higher GC in high temperature environment, which may be associated with reduced width of *E. coli* nucleoid in such conditions. Furthermore, our results also indicate that the GC of *E. coli* chromosome in stationary phase was greater than that in logarithmic phase for both normal and high temperature conditions (more pronounced in the former condition), and meanwhile the length of nucleoid in stationary phase is smaller under normal temperature and larger under high temperature compared to that in the logarithmic phase. Considering all these results together, there is no simple relationship between the chromosome GC and the nucleoid size in *E. coli* cells, meaning that GC is not a simple reflection of nucleoid size but an indicator of the intrinsic property of chromosome.

**Figure 4.**
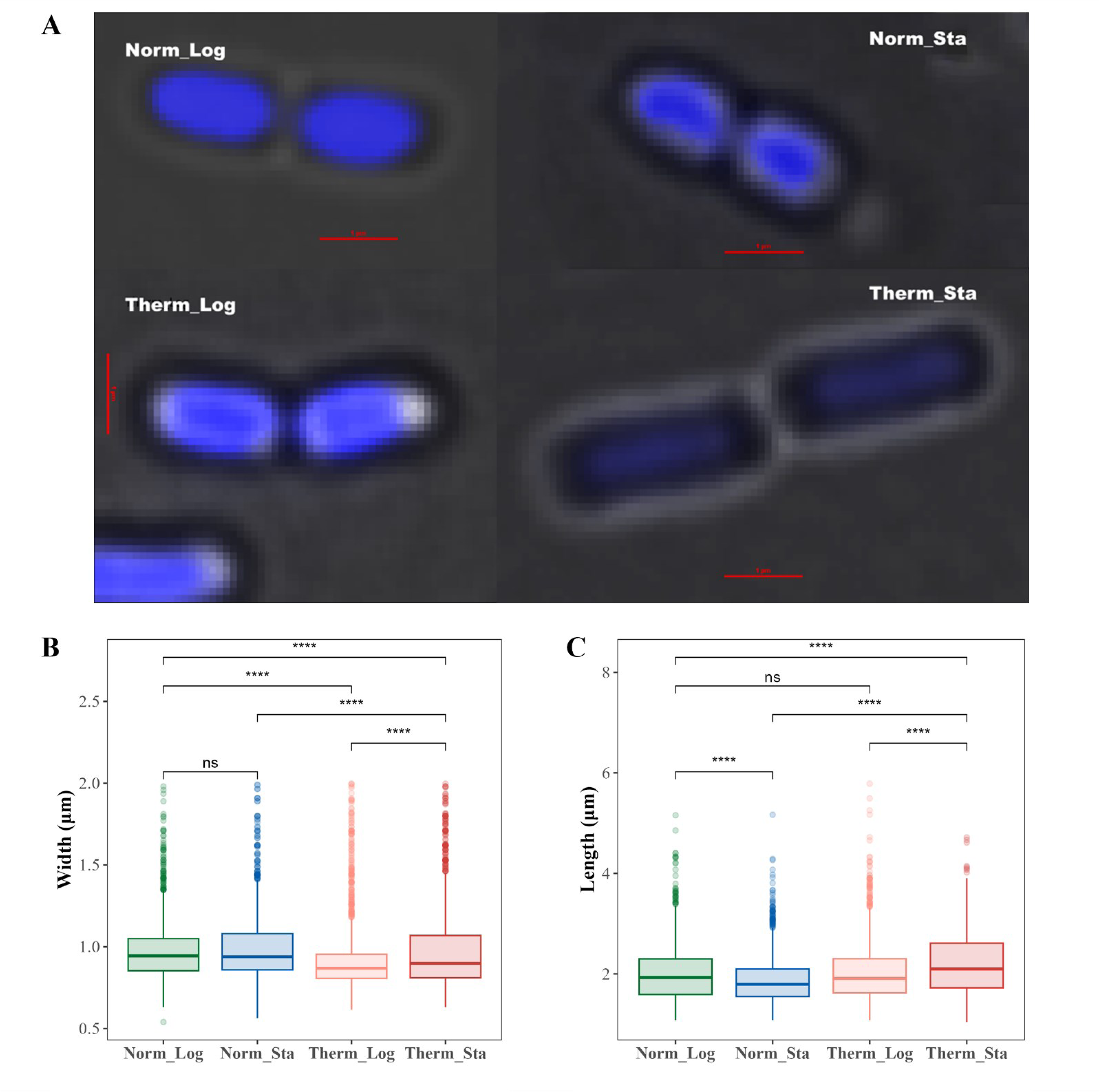
Comparison of *E. coli* nucleoid morphology in different growth conditions. (A) Typical microscopic images of *E. coli* nucleoid under various growth conditions. The nucleoids of *E. coli* were stained blue by DAPI. (B) Statistical results of *E. coli* nucleoid width under different growth conditions. (C) Statistical results of *E. coli* nucleoid length under different growth conditions. In B and C, significance levels are denoted by symbols: ns for *p* > 0.05, * for *p* < 0.05, ** for *p* ≤ 0 .01, *** for *p* ≤ 0 .001, and **** for *p* ≤ 0 .0001.

### 3.8. The Local Compactness of *E. coli* chromosome is affected by growth condition

To further characterize the change of chromosome 3D structure, we defined a metric called Local Compactness (LC). LC is calculated at various scales (up to 1000 kb) across the entire genome. **Figure 5A** depicts the heatmap representing the genome-wide LC for all analyzed scales. As the calculation scale increases, the overall LC increases, with a larger fluctuation observed at smaller scales. Notably, LC exhibits uneven distribution throughout the genome at any scale, with the Ter macrodomain displaying significantly lower compactness compared to other regions. This pattern persists across all growth conditions (heat stress or stationary phase). To evaluate the influence of growth condition on LC, we calculated the logarithmic ratio of LC between heat stress and normal temperature (**Figure 5A**, lower panels). At smaller scales (less than 50 kb), LC fluctuates considerably. However, at larger scales (more than 200 kb), the LC of the Ter macrodomain and its flanking regions significantly increases under high temperature, while the LC of the Ori macrodomain and its flanks decreases, and this decrease is more pronounced in the stationary phase.

**Figure 5.**
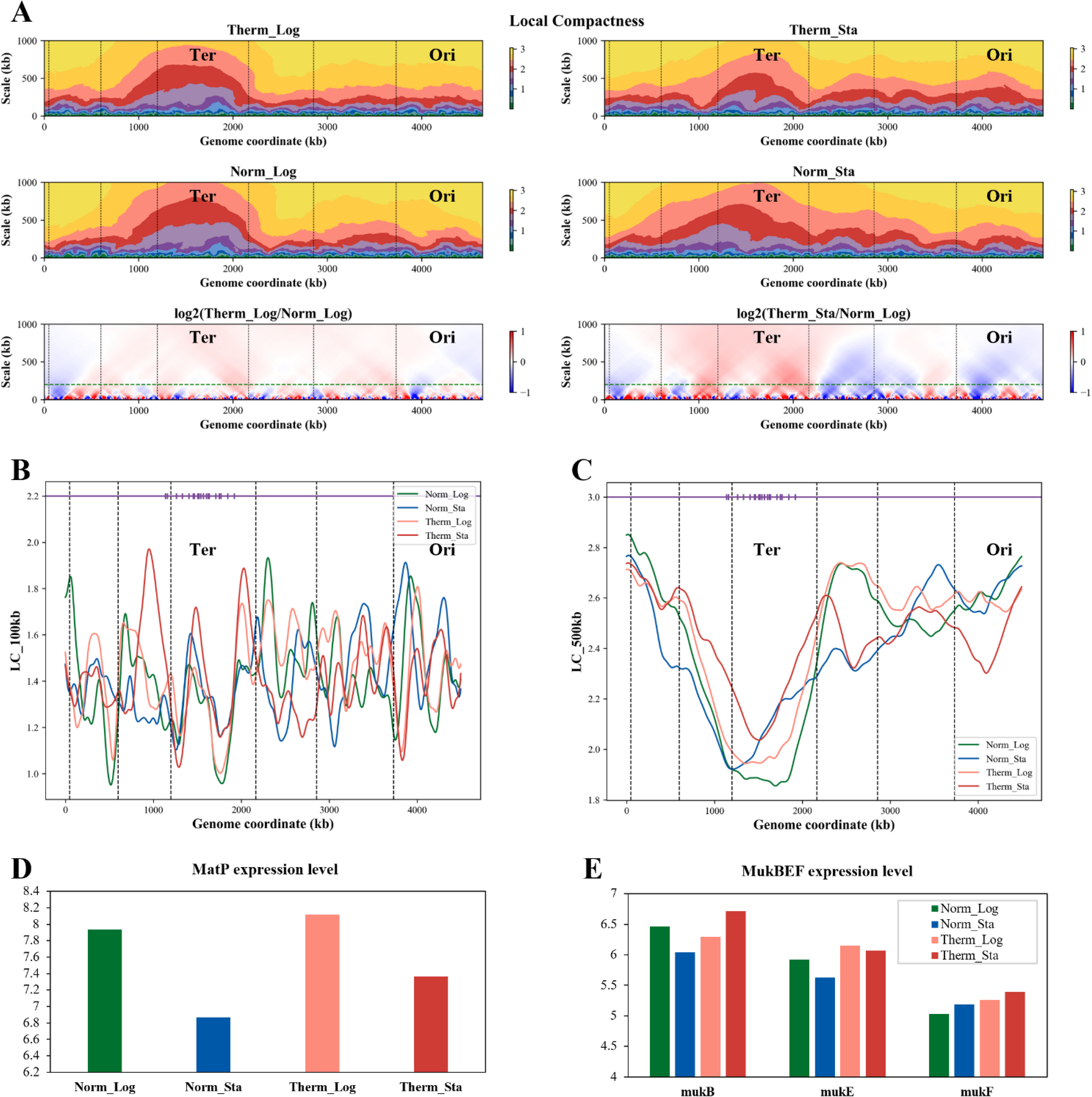
Local Compactness of the *E. coli* chromosome and related features. (A) The Local Compactness of *E. coli* chromosome 3D structures under different growth conditions and its ratio between high and normal temperatures. The green dashed lines in the bottom two subfigures of figure A indicate the scale of 200kb. (B) The Local Compactness in 100 kb range of the *E. coli* chromosome 3D structures under different growth conditions. (C) The Local Compactness in 500 kb range of the *E. coli* chromosome 3D structures under different growth conditions. In subfigures B and C, the lines of different colors correspond to different growth conditions. The dots on the purple horizontal lines represent the location of MatS which is the binding site of MatP. The black dashed vertical lines in the subfigure A, B, C represent the boundaries of macrodomains. (D) and (E) Transcription levels of MatP and MukBEF in *E. coli* under different growth conditions, respectively.

We further analyzed LC at two specific scales (100 kb and 500 kb) across all the four growth conditions, with the MatS location highlighted in purple (**Figure 5BC**). The 100 kb LC profiles display significant fluctuations, with less prominent differences observed at the Ter macrodomain boundaries. However, the 500 kb profiles reveal substantial LC increase at the Ter macrodomain boundaries under heat stress. This suggests a potential for increased interaction between the Ter macrodomain and distant DNA segments under high temperature conditions, consisting with the long-range interaction frequency analysis above. MatP, a key NAP, plays a crucial role in Ter macrodomain organization by specifically recognizing and binding to MatS sites on DNA. As shown in **Figure 5D**, MatP expression is upregulated under high temperature stress. Conversely, the MukBEF complex, involved in long-distance chromosome organization [11], exhibits varying degrees of upregulation under heat stress (**Figure 5E**). We hypothesize that high temperature stress alters MatP expression, leading to its enhanced binding to MatS sites and increased compactness of the Ter macrodomain. The MukBEF complex, on the other hand, might be more involved in mediating long-distance chromosome interactions under heat stress, potentially contributing to a reduction in the distances between the Ter macrodomain and other macrodomains (**Figure 3C**).

### 3.9. The Local Compactness is negatively correlated with transcription level

The Z-score curves illustrating the relationship between LC and transcription level are depicted in **Figure 6A**. We observed a negative correlation between LC and transcription level across the entire genome, with correlation coefficients being small but significant (*p* < 0.05). Heat stress triggers the accumulation of misfolded proteins in *E. coli*, and refolding or degradation of these aberrant proteins necessitates the involvement of numerous heat shock proteins (HSPs) [26]. Sigma factors, namely, σ^70^, σ^32^, σ^S^, and σ^E^, play a critical role in the transcriptional regulation of HSP genes. σ^70^ is encoded by rpoD, σ^32^ by rpoH, σ^S^ by rpoS, and σ^E^ by rpoE. We analyzed the LC changes in the chromosome regions harboring the genes (rpoD, rpoH, rpoE, and rpoS) encoding these sigma factors within the 3D models of *E. coli* chromosomes under various growth conditions (**Figure 6B**). These analyses revealed a strong negative correlation between LC and the corresponding transcription levels of the sigma factor genes. Overall, the correlation between LC and transcription level is negative and weak, but sometimes strong for specific genes.

**Figure 6.**
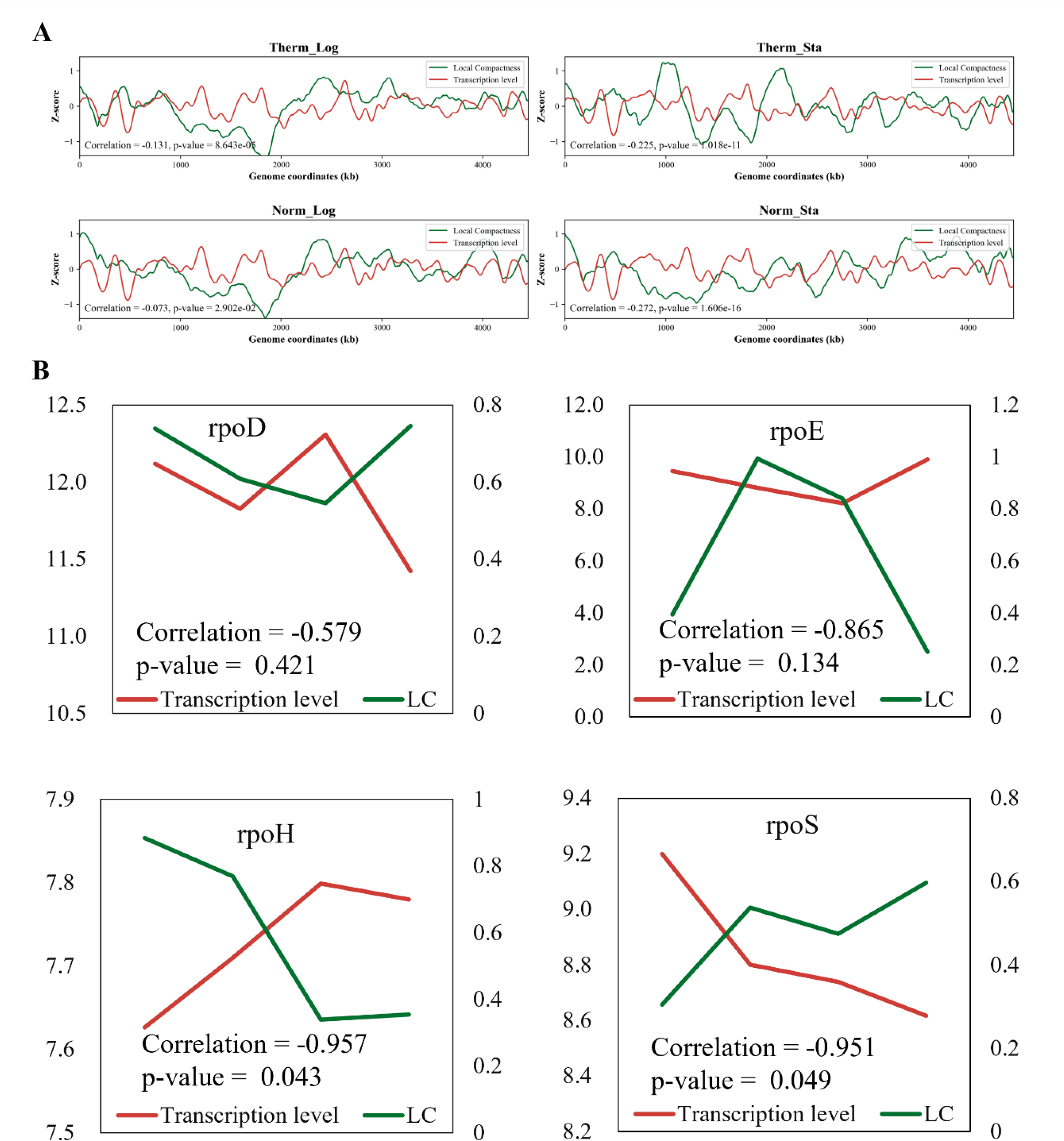
Correlation between Local Compactness and transcription level. (A) The genome-wide correlation between Local Compactness and transcription level under different growth conditions. (B) The correlation between the transcription levels of four sigma factors and their local compactness (where the sigma factor genes reside in the linear genome). Green lines correspond to local compactness; red lines correspond to transcription level.

## 4. Discussion

Bacteria constantly encounter environmental fluctuations, including temperature, pH, salinity, and others. These changes trigger stress responses, impacting their internal physiology and biochemistry, including transcriptional expression. These cellular alterations can directly or indirectly influence chromosome structure. 3C technology and its derivatives provide valuable tools to resolve the spatial relationships between DNA segments on chromosomes. This study successfully employed 3C-seq to capture the chromosome interaction data and built the 3D structure models of *E. coli* chromosomes. Furthermore, two metrics of GC and LC are defined for the 3D structural model to elucidate changes in chromosome conformation. It was observed that under high temperature stress, short-range interaction frequency of chromosome decreased while the GC of chromosome increased. The long-range interaction frequency of the Ter macrodomain increased, while its LC decreased. Correlations were found between transcription level and short-range interaction frequency. These findings shed light on the impact of heat stress on the structure and function of *E. coli* chromosomes.

NAPs play a crucial role in *E. coli* chromosome architecture and gene expression regulation. Environmental stress can affect the expression of NAPs. **Figure S4** depicts the transcription levels of key NAPs (HU, Fis, CbpA, H-NS, and StpA) under various growth conditions. CbpA, a non-sequence-specific DNA-bending protein [40], is negatively regulated by Fis during rapid growth [41]. Fis is known to mediate interactions between DNA segments exceeding 100 kb, HU is thought to facilitate interactions in the range of 50-280 kb, and H-NS is involved in long-range interactions (>250 kb) [9]. StpA is a kind of H-NS-like NAP that can form rigid DNA filaments with the function of transcription suppressor, and its non-specific binding affinity is 5 times higher than H-NS [42–45]. Thus, the observed decrease in short-range interaction frequency under heat stress may be attributed to the up-regulation of Fis and StpA transcription levels, and the decrease of CbpA and HU transcription levels.

In conclusion, the joint analysis of 3D genomics and transcriptomics revealed the impact of heat stress on the structure and function of *E. coli* chromosomes, which is crucial for understanding the molecular mechanism of 3D genome structure and heat adaptation in *E. coli*. The GC and LC metrics proposed in this work provide valuable tools for measuring and comparing chromosome 3D structures in future studies.

## Supporting information

Supplementary materials, methods and results, a PDF file.

## Acknowledgements

This work was supported by the National Natural Science Foundation of China (Grant 31971184).

