## Supplementary Information for "Spatial chromosome organization and adaptation of *Escherichia coli* under heat stress"

### 1. Supplementary materials

**Table S1. Regents**

| **Regent** | **Source** | **Identifier** |
|---|---|---|
| HpaII | New England Biolabs | Cat# R0171L |
| T4 DNA Ligase | New England Biolabs | Cat# M0202M |
| Ready-Lyse™ Lysozyme Solution | Epicentre | Cat# R1802M |
| Protease Inhibitor Cocktail | Sigma Aldrich | Cat# P8465 |
| Proteinase K | Transgene | Cat# GE201-01 |
| Glycogen | Thermo Fisher | Cat# R0561 |
| RNase A | Thermo Fisher | Cat# EN0531 |
| Triton™ X-100 | Sigma Aldrich | Cat# T9284 |
| DAPI Solution | Solarbio | Cat# C0065 |

**Table S2. Software**

| Software | Source |
|---|---|
| Bowtie2 | https://bowtie-bio.sourceforge.net/bowtie2/ |
| Python 3.7.1 | https://www.python.org |
| PyMol | https://pymol.org/2/ |
| Rstudio | https://www.rstudio.com/ |
| Cutadapt3 | https://pypi.org/project/cutadapt/ |
| Cluster Profiler | BiocManager |
| OpenCV | https://opencv.org/ |
| FastQC | https://www.bioinformatics.babraham.ac.uk/projects/fastqc/ |
| Tophat | http://tophat.cbcb.umd.edu/ |
| Cufflinks | http://cufflinks.cbcb.umd.edu/ |

### 2. Supplementary methods

#### 2.1 3C and RNA library construction

The method described in Lioy and co-authors' work [Cell 2018, 172: 771-783] was modified to generate a 3C library of *E. coli*. The process is described as follows. Fresh formaldehyde solution with a final concentration of 7% was added to 1mL of *E. coli* culture solution, incubated at room temperature for 30 min and then incubated at 4℃ for 30 min to fix chromosomes. The excess formaldehyde was quenched by adding 2.5 M glycine solution and incubating at 4 ℃ for 30 min. Centrifuge at 4 ℃ at 3500 × g for 10 min and remove the supernatant. 10 μL protease inhibitor and 200 μL Ready-Lyse were added to each tube of precipitation, and the precipitation was resuspended and incubated at room temperature for 30 min. Centrifuge at 4℃ at 3500 × g for 10 min and remove the supernatant. The bacterial lysate was obtained by re-suspension of precipitate products with 100 μL 1 × TE Buffer and incubation with 5 μL 10% SDS at room temperature for 20 min. Each 100 μL lysis solution was mixed with 50 μL 20% TritonX-100, 100 μL Cut Smart Buffer, 710 μL ultra-pure water and 40 μL restriction enzyme HpaII. After incubating at 37℃ for 6 h, the DNA was fully broken and 50 μL 10% SDS was added to quench the excess restriction enzyme. To obtain the ligase, 100 μL

BSA, 1 mL 20% TritonX-100, 2 mL T4 ligation Buffer, 1.6 mL ultra-pure water, and 10 μL T4 DNA ligase were added to each 1 mL enzyme digestion solution. After incubating at 16°C for 4 hours, add 200 μL 0.5 M EDTA and mix well. Proteinase K was added to the mixed solution at a final concentration of 100 μg/mL. Incubated at 58°C overnight. The 3C library was obtained by extracting DNA from the above mixed solution.

RNA library construction was conducted by Novogene company, and the protocol is as follows. RNA was harvested using Rneasy mini plus kit (Qiagen). 1.3 ug of total RNA was used for the construction of sequencing libraries. After the RNA samples passed the detection, the mRNA was enriched by removing rRNA through the kit. Then mRNA was fragmented into fragments by fragmentation buffer. A strand of cDNA was synthesized by using mRNA as template. Then buffer, dNTPs (dTTP in dNTP was replaced by dUTP) and DNA Polymerase I and RNase H were added to synthesize the double-stranded cDNA. AMPure XP Beads were used to purify the double-stranded cDNA, and USER enzyme was used to degrade the second strand of cDNA containing U. The purified double-stranded cDNA was repaired at the end, A tail was added, and sequencing adaptor was connected. Then AMPure XP beads were used for fragment size selection. Finally, PCR amplification was performed and AMPure XP beads were used to purify the PCR products to obtain the final library. After the construction of the library, Qubit 2.0 was used for preliminary quantification, and then Agilent 2100 was used to detect the size of the inserted fragments of the library. After the inserted fragments met the expectations, q-PCR method was used to accurately quantify the effective concentration of the library to ensure the quality of the library.

#### 2.2 Calculations of short-range interaction frequency and proportion

The short-range interaction frequency is obtained by dividing the sum of the interaction frequencies between the target bin and two bins with the same distance (threshold) from the target bin by the sum of the interaction frequencies between the target bin and all bins. The formula is as follows:

$$sf_i = \frac{f(i, i - thr) + f(i, i + thr)}{\sum_{j=0}^{n} f(i,j)}$$

In the formula, $sf_i$ is the short-range interaction frequency of bin $i$, and $thr$ is the threshold value to define the short range. In this work, $sf$ values for the thresholds from 0 to 200 bins were calculated.

The short-range interaction proportion is obtained by dividing the sum of the interaction frequencies of the target bin with all bins within a short range (defined by a threshold) by the sum of the interaction frequencies of the target bin with all bins in the genome. The formula is as follows:

$$sp_i = \frac{\sum_{j=i-thr}^{i+thr} f(i,j)}{\sum_{j=0}^{n} f(i,j)}$$

In the formula, $sp_i$ is the short-range interaction proportion of bin $i$, and *thr* is the threshold value to define the short range. In this work, *sp* values for the thresholds from 0 to 200 bins were calculated.

### 2.3 Global Compactness calculation

Since bacterial chromosome is usually in a ring shape, we consider its loosest state is a regular circle. When the generated 3D model of a chromosome is closer to a regular circle, the chromosome is considered less compact. The overall structure is least compact when it forms a circle, and the structure becomes more compact when any part deviates from the circle. **Figure S1** illustrates the change of compactness when the shape deviates from circle. As shown, a regular circle has a compactness of 0, which is the loosest state. The compactness of a hexagon is 0.084, the compactness of a circle with a right angle is 0.248, and the compactness of a circle with two symmetrical right angles is 0.509.

To calculate the global compactness of a bacterial chromosome, first we need to calculate the circumference of the chromosome. With the spatial coordinates of each bin in the 3D structure model, we can calculate the Euclidean spatial distance *D* between any two bins. The sum of the distances between all successive bins is regarded as the total length of the chromosome model *L*. The formula for *L* is as follows:

$$L = D(0,n) + \sum_{i=1}^{n} D(i,i-1)$$

In this formula, *L* is the length of the chromosome model, *n* is the total number of bins in the chromosome model, and *D* is the Euclidean distance between bins.

A regular circle model is then reconstructed using the length *L* of the chromosome model as the perimeter. Calculating the distance between bins on the reconstructed circle requires first arranging this reconstructed circle on a rectangular coordinate system. According to the circumference formula of a circle $r = L/(2\pi)$, the radius *r* of the reconstructed circle can be calculated. The starting bin of the chromosome model is placed on the *x*-axis of the coordinate system, and the center of the reconstructed circle is placed at the origin point of the coordinate system; then, the coordinates of the starting bin are (*r,* 0), and the coordinates of the rest bins are computed according to their relationship with the first bin. When the chromosome is reconstructed into a circle, the spatial distance between two successive bins is transformed into the arc length *l* on the reconstructed circle. The arc length $l$ of a target bin $m$ from the starting bin on the reconstructed circle can be calculated as follows:

$$l(0, m) = \sum_{i=1}^{m} D(i, i-1)$$

In this formula, $l$ (0, $m$) is the arc length between the target bin $m$ and the starting bin 0, and $D$ is the Euclidean spatial distance between two successive bins.

Next, according to the formula $rad(0, m) = l(0, m)/r$, calculate the corresponding radian *rad*. Then the coordinates $x$ and $y$ of the target bin $m$ in the rectangular coordinate system are calculated according to the sine and cosine formulas $x_m = r * \cos\big(rad(0, m)\big)$ and $y_m = r * \sin\big(rad(0, m)\big)$. According to the Euclidean distance calculation method, the distances $d$ $(i, j)$ between all bin pairs $i$ and $j$ on the reconstructed circle can be obtained. And then the Global Compactness of chromosome structure model can be calculated as follows:

$$GC = -\log_2 \frac{\sum_{i,j=0,i\neq j}^{n} D(i, j)}{\sum_{i,j=0,i\neq j}^{n} d(i, j)}$$

In this formula, GC is the Global Compactness, $i$ and $j$ are indexes for bins, $D$ $(i, j)$ is the Euclidean distance between bin $i$ and bin $j$ in the chromosome 3D structure model, and $d$ $(i, j)$ is the Euclidean distance between bin $i$ and bin $j$ calculated based on their new coordinates $(x_i, y_i)$ and $(x_j, y_j)$ when the bins are rearranged on the reconstructed circle. The sum of Euclidean distances between all bins in the chromosome 3D structure model and the sum of Euclidean distances between all bins on the reconstructed circle were calculated, and then the negative log2 value of the ratio of the two sums was defined as the Global Compactness of chromosome. The larger the value, the higher the compactness. **Figure S2A and B** show the Global Compactness of six theoretical models.

**Schematic diagram of compactness calculation**

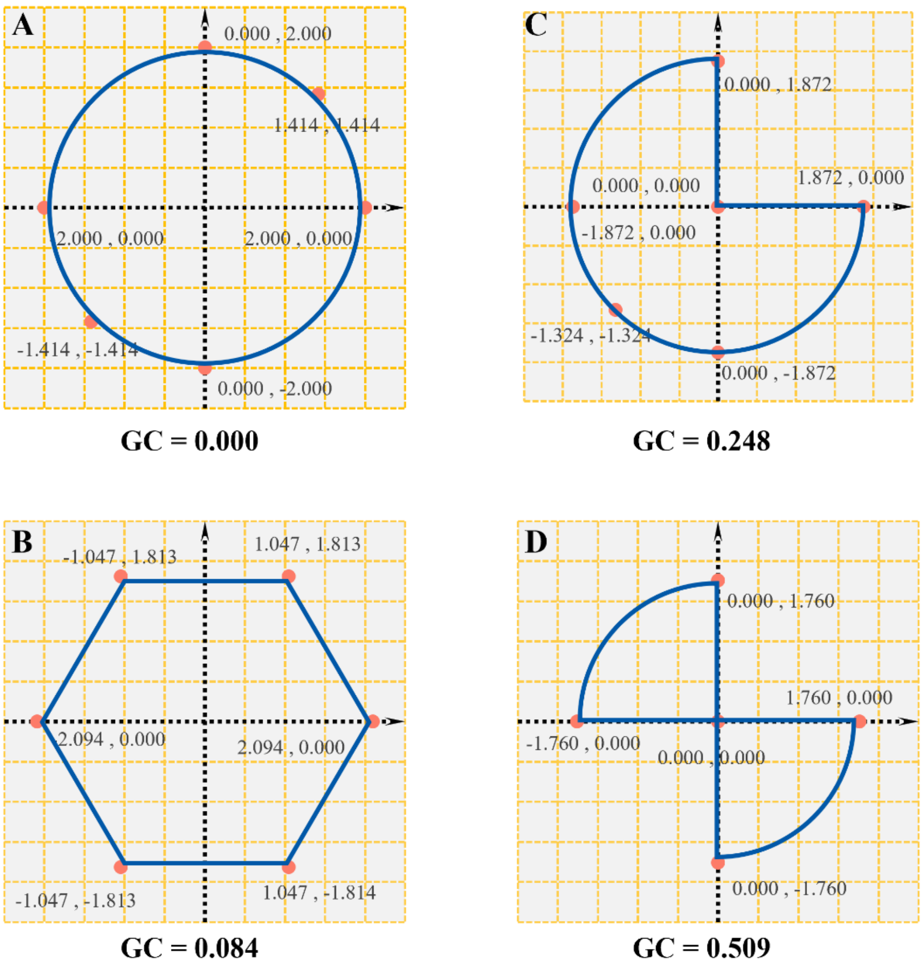


**Figure S1. Global Compactness calculation diagram.** (A) Suppose a circular coil with radius 2 has a global compactness of 0. (B) Keeping perimeter the same, when this coil is reshaped into a regular hexagon, its radius becomes 2.094 and the global compactness is 0.084. (C) Keeping perimeter the same, if the coil is retracted by a right angle to the center of the circle, the radius becomes 1.872 and the global compactness is 0.248. (D) Keeping perimeter the same, if the coil is reshaped into two symmetrical right angles towards the center of the circle, the radius becomes 1.760 and the global compactness is 0.509.

### 2.4 Local Compactness calculation

The local chromosome structure can be regarded as a line of bins, and the compactness is minimum when it is a straight line. The closer the local chromosome structure is to a straight line, the lower the compactness of that local position. Any deviation from the straight line makes the structure more compact. First, the bins within a certain range of the target bin (upstream and downstream symmetrically) are placed on a reconstructed straight line, and the length $L$ of the straight line is calculated as the sum of the Euclidean spatial distances between all successive bins within that range. Then the ratio of the sum of the Euclidean distances between these bins in the chromosome 3D structure model to the sum of the distances between these bins on the reconstructed straight line is calculated, and the negative log2 value of this ratio is taken as the local compactness value. The formula for Local Compactness is as follows:

$$LC_i = -\log_2 \frac{\sum_{j=i-thr}^{i+thr} D(i,j)}{\sum_{j=i-thr}^{i+thr} d(i,j)}$$

In this formula, $LC_i$ is the compactness in the local range of bin $i$, $D(i, j)$ is the Euclidean spatial distance between $i$ and $j$ in the chromosome model, $d(i, j)$ is the Euclidean distance between bin $i$ and bin $j$ on the reconstructed straight line in the local range, and *thr* is the threshold for local range definition. The larger the *LC*, the higher the chromosome compactness at that location. **Figure S2C** shows the Local Compactness of six theoretical models.

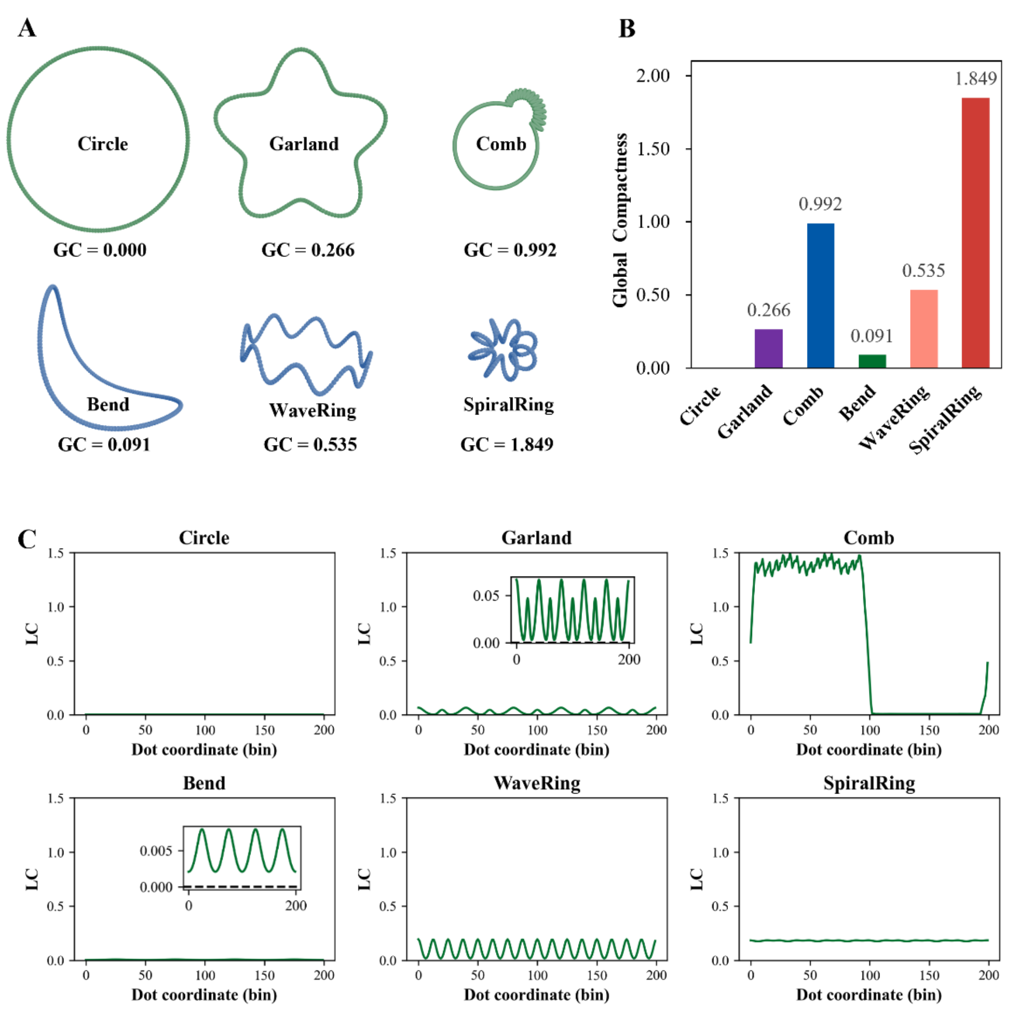


**Figure S2. Global Compactness and Local Compactness of six theoretical models.** (A) In different shapes, 200 points are connected into a ring. The three rings in the first row are planar (2D) structures, and the three rings in the second row are 3D structures. (B) Global Compactness of the six theoretical models. (C) Local Compactness of the six theoretical models.

### 3. Supplementary results

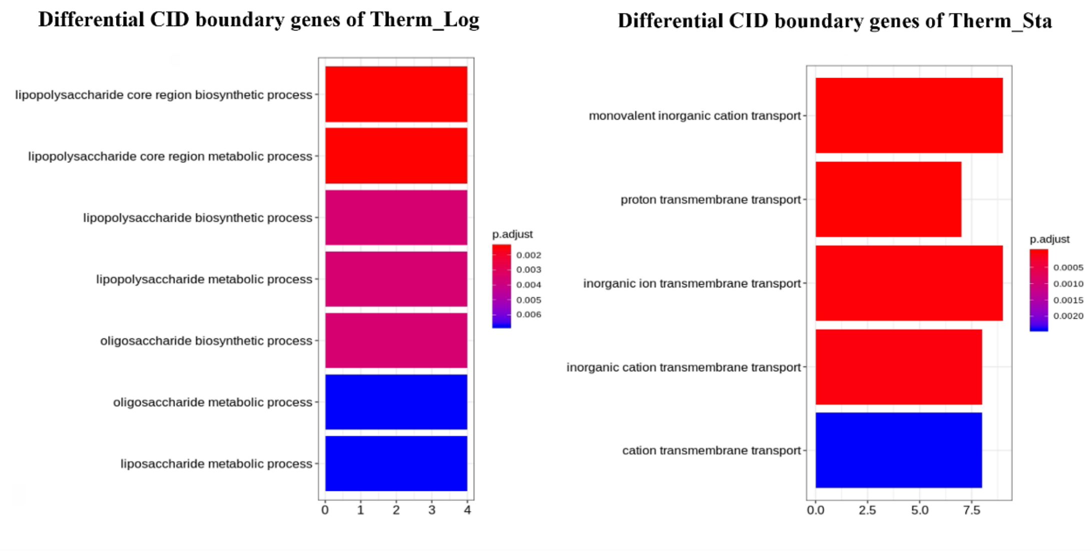


**Figure S3. GO enrichment results of chromosome differential CID boundary genes in *E. coli* under heat stress.** The left side is the result of high temperature logarithmic phase, and the right side is the result of high temperature stationary phase.

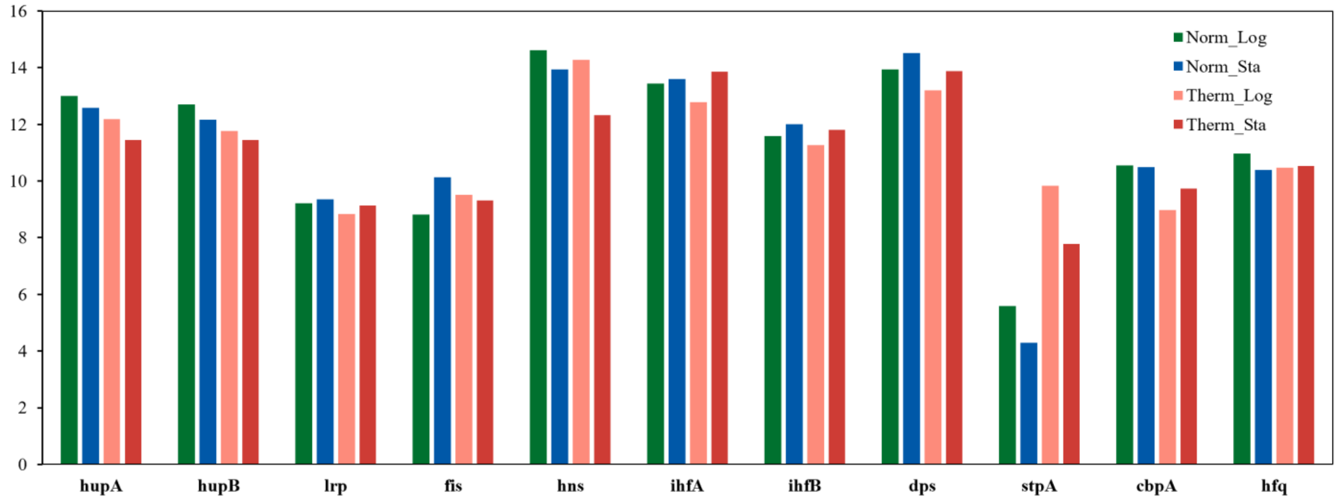


**Figure S4. Transcription levels of some *E. coli* NAPs under different growth conditions.** Different colors in the figure represent different growing conditions.
